# Adaptive decision-making by ants in response to past, imminent, and predicted adversity

**DOI:** 10.1101/2024.12.17.628737

**Authors:** Yusuke Notomi, Shigeto Dobata, Tomoki Kazawa, So Maezawa, Shigehiro Namiki, Ryohei Kanzaki, Stephan Shuichi Haupt

**Author notes:** Corresponding author, Stephan Shuichi HAUPT. **CONTACT INFORMATION** Yusuke NOTOMI.

## Abstract

Many animals exhibit innate behaviours, which are often interpreted as hard-wired, reflex-like responses, particularly in insects. Among these behaviours, beacon-aiming—an approach towards dark areas or objects—is observed in many animals; however, its functional significance remains unclear, and some ant species do not exhibit it. Here we show that in one such species, *Camponotus japonicus,* the behaviour was triggered only under adverse substrate conditions, such as during the crossing of liquid-covered surfaces, regardless of the locomotor patterns like walking or swimming, or the mere presence of water, or when walking upside-down. Once initiated, beacon-aiming persists even under normal substrate conditions, as demonstrated by ants transitioning from water-covered to dry substrates suitable for comfortable walking. This behavioural flexibility indicates that the innate behaviour is not hard-wired but modulated by internal states. Furthermore, changes in internal states may serve adaptive decision-making, potentially allowing ants to prepare for future adverse conditions. The isolated ants on a water-surrounded platform gradually established an attraction to the direction of the beacon before ultimately swimming towards it. These findings suggest that beacon-aiming is regulated by internal states, especially anxiety-like states formed in response to past, imminent, and predicted adverse substrate conditions in ants.

## INTRODUCTION

In dynamically changing environments in conjunction with varying internal conditions, animals must employ flexible decision-making to ensure survival. Several examples of remarkable cognitive processing in animals are known to significantly contribute to this goal ^1,2^. Importantly, cognitive functions are not strictly dependent on brain size ^3^. While larger brains facilitate advanced cognitive processing, their development and operation entail significant metabolic costs ^4,5^. Even with small brains ^6,7^, insects exhibit behaviours that involve cognitive processing^8,9^, but also strongly rely on simple stimulus-dependent responses, including innate behaviours, which have low processing requirements. Innate behaviours are considered to offer the advantage of rapid responses; however, they are still generally thought to be less flexible ^10^.

Inflexible behaviours can lead to disadvantages ^11^ so it makes sense that innate behaviours can be modified through learning ^12–14^. Previous studies have revealed that innate behaviours can be temporarily modulated by physiological factors ^15,16^. Nonetheless, the neural mechanisms underlying the dynamic modulation of innate behaviours remain poorly understood ^17,18^. This gap partly arises from the paucity of simple experimental paradigms to manipulate decision-making processes involved in innate behaviours, which are defined by their stability and induction through easily controlled stimuli ^19,20^.

Here, we demonstrate that an innate behaviour in an ant is controlled by internal states and present a simple paradigm in which internal state dynamics can be visualised through the behaviours. We also show that adaptive decisions taking into account future conditions— mainly associated with advanced cognitive processing, such as inference—does not necessitate learning.

The innate behaviour focused on in this study is ‘beacon-aiming’ ^21^ consisting of a spontaneous orientation towards visually conspicuous dark areas or objects. This behaviour has been observed across almost all insect orders ^22^, other invertebrates ^23,24^, and some vertebrates ^23^, and has also been reported in insects that fall onto water surfaces ^25,26^. In ants, beacon-aiming on land or water has been documented in various species ^27,28^ and is typically interpreted as an innate response, independent of age, feeding state, or caste ^22,29,30^. However, a comparative study found that some ant species do not exhibit beacon-aiming ^22^.

In the present study, we departed from the hypothesis that adverse substrate conditions could be a trigger for beacon-aiming. Choosing a species that did not perform the innate behaviour in previous tests, the partially arboreal ants *Camponotus japonicus* ^31^, we show that it does perform beacon-aiming when substrate conditions are adverse, such as immersion in water or walking upside-down. We then introduce experimental paradigms in which an ant has to transition from one substrate to another, i.e. from water to dry substrate and vice-versa. This set-up allows us to probe the decision-making dynamics in ants before, during, and after transition between substrates.

## RESULTS

### Experiment (i): Factors influencing approach behaviour towards a visual landmark

Orientation towards landmarks has been studied in different species using different environmental conditions such as substrates, but a comparison of orientation under different conditions in any species appears to be lacking. To elucidate the mechanisms underlying the emergence and regulation of context-dependent behavioural differences, we investigated whether workers of *C. japonicus* orient differently on land and water surfaces. Initially, two experimental conditions were prepared: ‘Land-walking’ – walking on a dry substrate (Figure 1A); and ‘Water-swimming’ – forced swimming across a water surface (Figure 1B) with a landmark (‘beacon’) placed on the arena wall. To exclude the possibility that the observed behaviour is specifically associated with locomotion unique to swimming, we also provided the ants with a ‘Water-wading’ condition where a shallow water depth allowed the ants to walk (wade) using a walking locomotor pattern (Figure 1B). The direction of the vector connecting the centre of the arena to the position where the ant reached a specified arena edge was defined as the final bearing of each trial.

**Figure 1.**
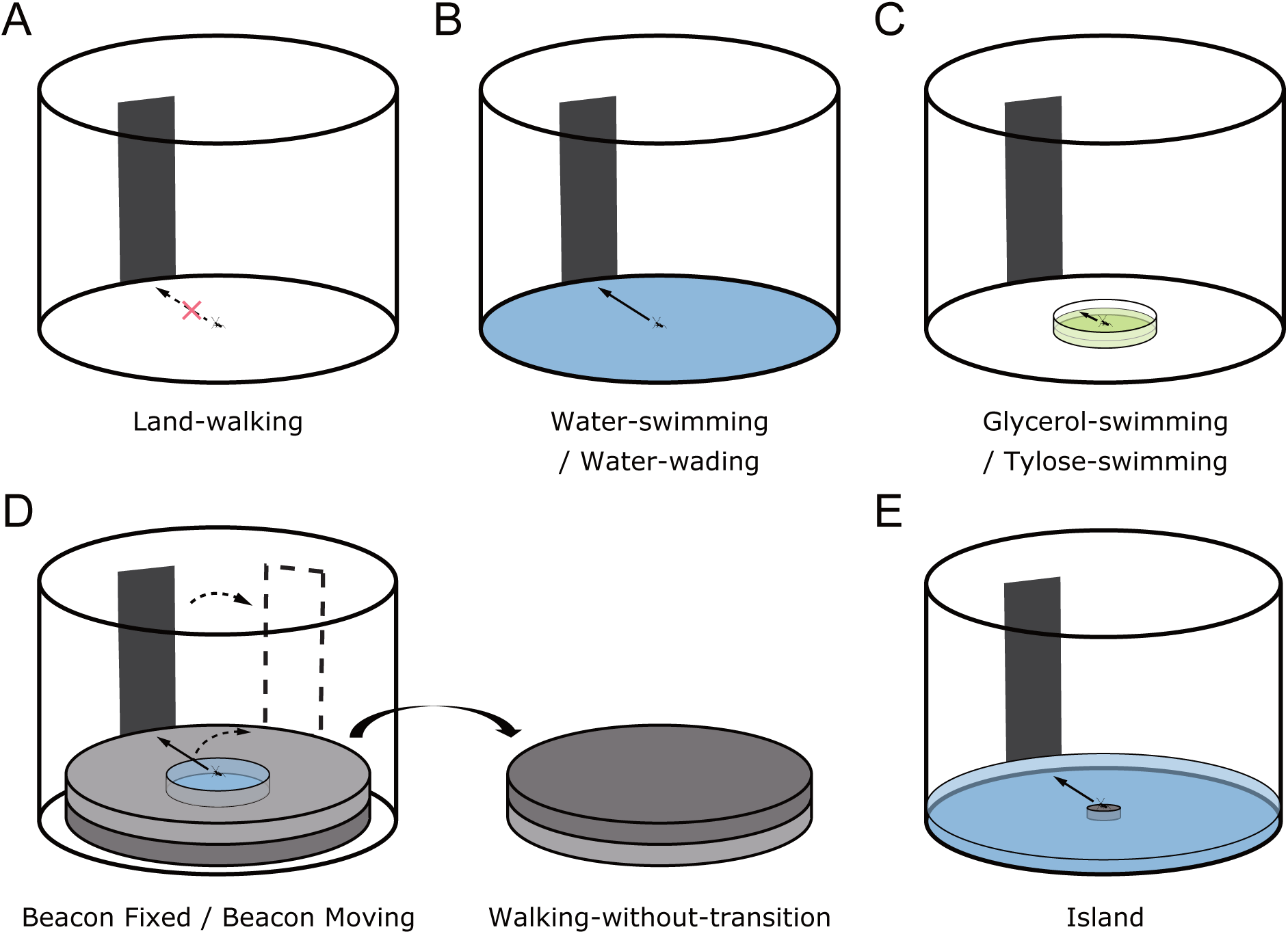
Experimental setup. All experiments were conducted in an arena with a black beacon (height: 45°, width: 20°, measured from the effective centre of the floor) placed against the wall. Arrows and cross marks represent the overall orientation trend. (A) Basic arena used for Land-walking. (B) Arena filled with water for swimming (Water-swimming) or with shallow water for wading (Water-wading). (C) Arena, same as (A), with a transparent dish filled with either a 80% (v/v) glycerol and 20% (v/v) 2-propanol mixture (Glycerol-swimming) or a 1.0% (w/w) tylose aqueous solution (Tylose-swimming). (D) Arena fitted with extruded polystyrene foam insets to enable the transition from swimming in water to walking on a dry substrate. One side of the inset contained a central pool that could be filled with water (for the conditions Beacon Fixed, Beacon Displaced), while the other side served as a control (for condition Walking-without-transition), providing only the uniform dry substrate. The final position of the beacon, which was displaced by 60° under the Beacon Displaced condition, is also illustrated (dashed outline). (E) Arena, same as (B), with a transparent cup in the centre and the water level matched to the base of the cup (Island), effectively creating a central platform surrounded by water (“island”).

Under Land-walking conditions, ant trajectories were relatively straight (Figure 2, Figure S1 in the supplemental information); however, their orientation appeared to be in random directions (Figure 2A). This was reflected by a wide interquartile range spanning more than 90 degrees in the circular boxplot of the final bearings, indicating no clear directional preference (Rayleigh test: *R̅* = 0.19, *P* = 0.140, *N* = 57).

**Figure 2.**
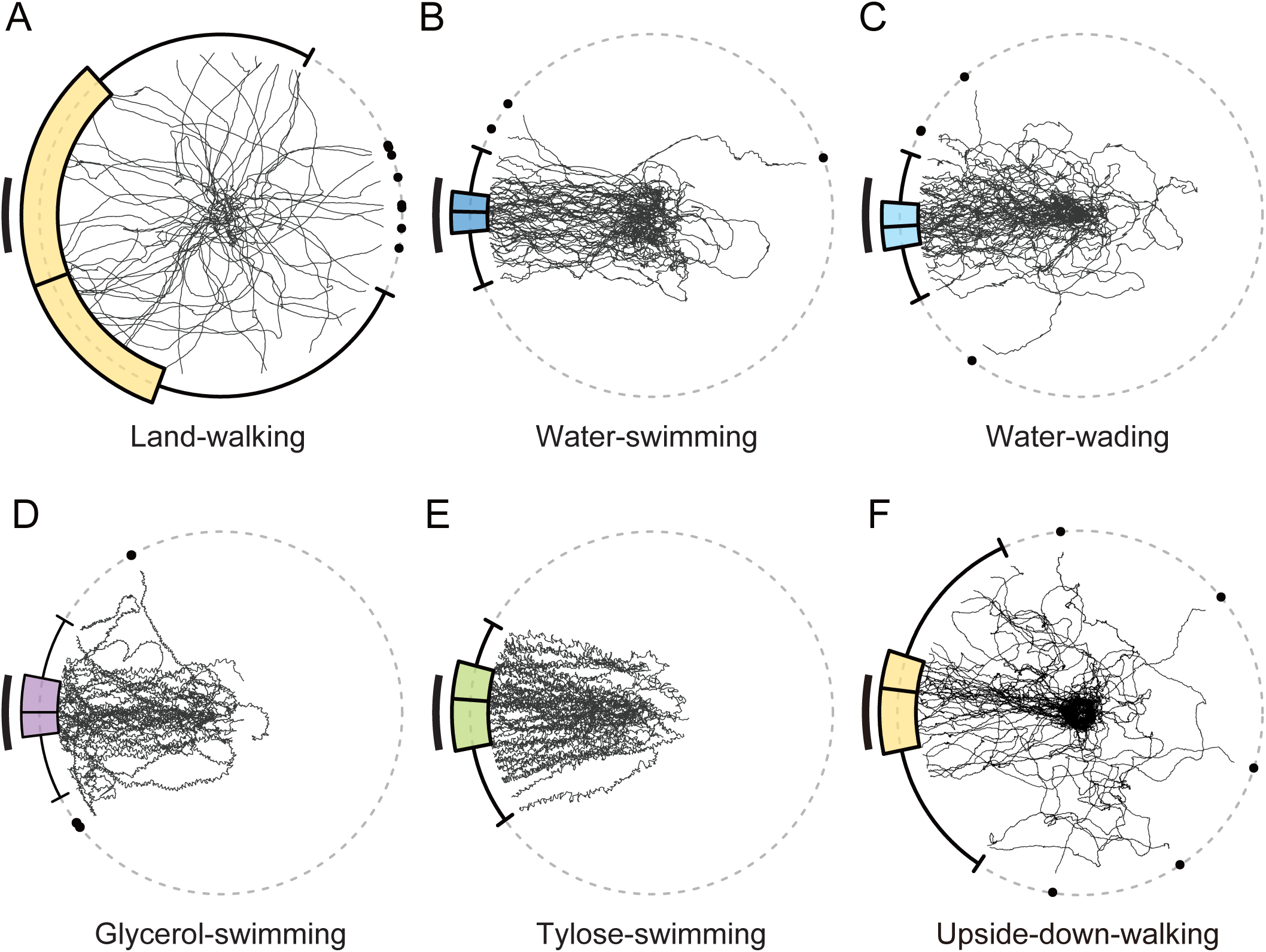
Beacon-aiming during walking on dry substrate, swimming, and wading in water. (A-F) Trajectories and circular box-and-whisker plots of final bearings are shown under the following conditions: (A) Land-walking (walking on dry substrate, *N* = 57); (B) Water-swimming (swimming in water, *N* = 57); (C) Water-wading (walking in shallow water, *N* = 57); (D) Glycerol-swimming (swimming in an 80% (v/v) glycerol and 20% (v/v) 2-propanol mixture, *N* = 47); and (E) Tylose-swimming (swimming in a 1.0% (w/w) tylose aqueous solution, *N* = 59); (F) Upside-down-walking (*N* = 45). The beacon range is represented as a black arc segment on the left side of each plot. Individual trajectories correspond to single trials. Circular box-and-whisker plots show the median, first quartile, third quartile (coloured interquartile range), minimum, maximum (whiskers), and outliers (black-filled circles) of the final bearing distributions.

Under Water-swimming conditions, except for occasional difficulties, ants adopted a distinctive swimming posture with their antennae raised above the water and a locomotor pattern characterised by extended hind legs. During swimming, most trajectories meandered slightly but generally oriented towards the beacon or its vicinity (Figure 2B). The interquartile range of the circular boxplot of the final bearings of Water-swimming conditions overlapped with the beacon range and was relatively narrow, indicating a preference for the centre of the beacon (Rayleigh test: *R̅* = 0.95, *P* < 0.001, *N* = 57; mean direction: 1.3°, 95% confidence interval: [-1.8°, 4.4°]). The final bearing distribution was significantly different from Land-walking (Mardia–Watson–Wheeler test: *W*_(2)_ = 73.01, *P* < 0.001).

Under Water-wading conditions, the distribution of final bearings (Rayleigh test: *R̅* = 0.97, *P* < 0.001, *N* = 57; mean direction: −2.1°, 95% confidence interval: [−5.9°, 1.6°], Figure 2C) was similar to that in Water-swimming. The final bearing distribution was significantly different from Land-walking (Mardia–Watson–Wheeler test: *W*_(2)_ = 69.28, *P* < 0.001) but not significantly different from Water-swimming (Mardia–Watson–Wheeler test: *W*_(2)_ = 3.12, *P* = 0.629). These results suggest that the induction of beacon-aiming in water is not triggered by locomotor patterns but is instead influenced by water itself in *C. japonicus*.

Furthermore, the involvement of water reception in beacon-aiming was assessed by replacing water with two kinds of liquids (Figure 1C). For ‘Glycerol-swimming’, ants swam in a mixture of glycerol and 2-propanol. For ‘Tylose-swimming’, ants swam in a tylose aqueous solution adjusted to closely match the dynamic viscosity of the glycerol-propanol mixture. When ants introduced into these liquids, the smooth paddling motion was hindered by the high viscosity of these liquids. Under ‘Glycerol-swimming’ conditions, they swam towards the beacon across the liquid surface, although their locomotion was impaired (Glycerol-swimming, Rayleigh test: *R̅* = 0.95, *P* < 0.001, *N* = 37; mean direction: 1.5°, confidence interval: [−4.2°, 6.9°], Figure 2D). Under ‘Tylose-swimming’ conditions, ants also swam towards the beacon (Rayleigh test: *R̅* = 0.96, *P* < 0.001, *N* = 49; mean direction: 1.0°, con_fidence interval: [−3.3°, 5.2°], Figure 2E). No significant difference was observed in the distribution of final bearings between swimming across these two liquids (Mardia–Watson–Wheeler test: Glycerol-swimming and Tylose-swimming: *W*_(2)_ = 2.66, *P* = 0.265). These results indicate that the presence of water is not required for the induction of beacon-aiming in the ants.

In addition, as an adverse substrate condition that does not employ liquids at all, we forced ants to walk upside-down (‘Upside-down-walking’) on sandpaper. Under this condition, 32/77 ants dropped from the substrate, demonstrating it is difficult to negotiate for them. This adverse substrate condition also induced their beacon-aiming (‘Upside-down-walking’, Rayleigh test: *R̅* = 0.72, *P* < 0.001, *N* = 45; mean direction: 10.0°, confidence interval: [−2.7°, 24.1°], Figure 2F). Therefore, liquid immersion is not necessary to induce beacon-aiming, but it is considerably more convenient in experimental situations.

### Experiment (ii): Persistence of beacon-aiming decision made after water-to-land transition

To investigate whether the context-dependent behavioural changes are direct sensory responses corresponding to environments or regulated in a more sustained manner, we next examined whether the directional decision persisted after a transition from water to a dry substrate (equivalent to aquathlon, Figure 1D). The experiment was conducted under three conditions: ‘Beacon Fixed’ – the beacon remained stationary on the arena wall with a central pool surrounded by walkable dry area (shore/land); ‘Beacon Displaced’ – the beacon was rapidly shifted by 60° upon the transition of the ants from water to dry substrate; and ‘Walking-without-transition’ – ants walked on a dry arena without a pool. The boundary between land and water was defined as the pool edge.

Swimming in a pool induced beacon-aiming, shown as the distributions of the final bearings on the pool edge under all conditions (Beacon Fixed, Beacon Displaced: Rayleigh test: *R̅* = 0.85, 0.84, both *P* < 0.001, *N* = 33, 33, mean direction: 1.0°, 0.0°, confidence interval: [−8.2°, 9.2°], [−7.9°, 7.3°], Figure 3B, C, respectively).

**Figure 3.**
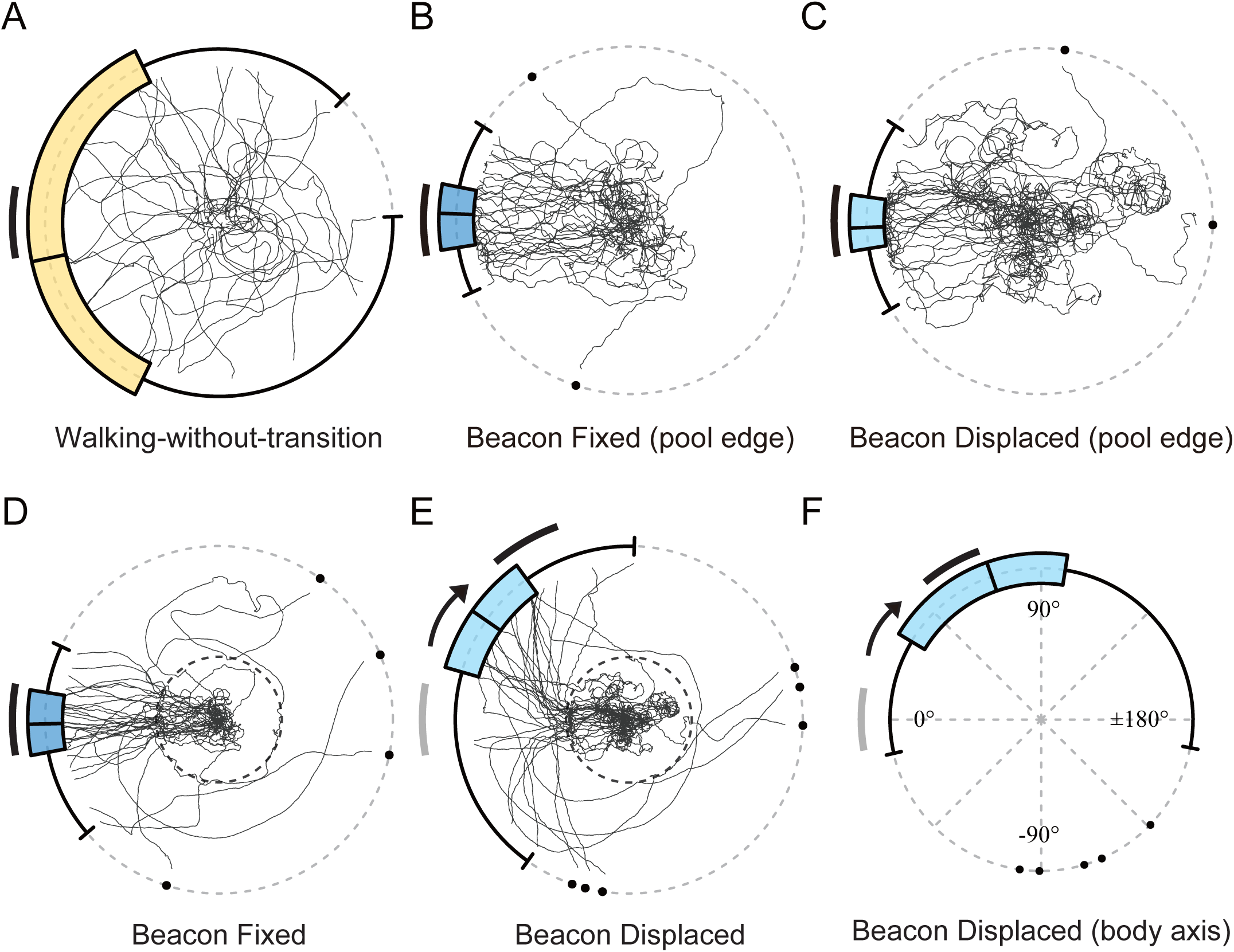
Persistence of beacon-aiming decision made after water-to-land transition. (A-E) Trajectories and circular box-and-whisker plots of final bearings are shown under the following conditions: Walking-without-transition condition, until ants reached the arena edge (A, *N* = 33); Beacon Fixed condition, until ants reached the pool edge (B, *N* = 33); Beacon Displaced condition, until ants reached the pool edge (C, *N* = 33); Beacon Fixed condition, until ants reached the arena edge (D, *N* = 33); and Beacon Displaced condition, until ants reached the arena edge (E, *N* = 33), following the format of Figure. 2. For the condition where the beacon was displaced by 60° (E), an arrow indicates the direction of the beacon shift. The initial beacon range is shown in grey, the final beacon range after displacement in black. (F) Circular box-and-whisker plots illustrate the distribution of the body axis direction (from abdomen to head) at the arena edge under Beacon Displaced condition, as described in (E).

Under Beacon Fixed conditions, beacon-aiming persisted after the transition from water to land (Rayleigh test: *R̅* = 0.78, *P* < 0.001, *N* = 33, mean direction: −2.7°, confidence interval: [−10.5°, 5.3°], Figure 3D). No significant difference was observed in the distributions of final bearings at the pool edge and arena edge (Mardia–Watson–Wheeler test: *W*_(2)_ = 0.27, *P* = 0.875).

Under Walking-without-transition conditions on the same dry polystyrene substrate, no orientation towards the beacon was observed (Rayleigh test: *R̅* = 0.28, *P* = 0.145, *N* = 33, Figure 3A, Figure S2 in the supplemental information). The distribution of final bearings at arena edge in the water to land transition experiment under Beacon Fixed conditions was significantly different from Walking-without-transition (Mardia–Watson–Wheeler test: *W*_(2)_ = 25.74, *P* < 0.001, *N* = 33, 33).

Under Beacon Displaced conditions, the beacon-aiming persisted and the directional decision was updated even after the transition from water to land, as indicated by the mean direction shifting towards the new beacon position, which was displaced by 60° (Rayleigh test: *R̅* = 0.61, *P* < 0.01, *N* = 33, mean direction: 31.5°, confidence interval: [12.1°, 48.6°], Figure 3E). This shift resulted in a significant difference between the distributions of final bearings at the pool edge and the arena edge (Watson-Wheeler test: *W*_(2)_ = 30.58, *P* < 0.001) as well as between the distributions of final bearings at arena edge under Beacon Fixed and Beacon Displaced conditions (Mardia–Watson–Wheeler test: *W*_(2)_ = 30.69, *P* < 0.001). Final bearings in the Beacon Displaced condition exhibited a somewhat variable partial shift to the new beacon position such that no significant difference was found in comparison to the Walking-without-transition condition (Mardia–Watson–Wheeler test: *W*_(2)_ = 7.72, *P* = 0.021, *N* = 33, 33, with the Bonferroni correction accounted for multiple comparisons among three conditions). However, the body axis direction was significantly oriented towards the new beacon position (Rayleigh test: *R̅* = 0.50, *P* < 0.001, mean direction: 68.6°, confidence interval: [45.1°, 92.7°], Figure 3F). Once the ants reached the shore, they did not re-enter the water.

The behaviour observed before and after the transition from water to land suggests that the beacon-aiming decision persists even after the ant has escaped the adverse substrate condition. Furthermore, their decision-making appears to be influenced not only by the current substrate condition but also by the immediately preceding adverse condition, while continuously updating their orientation. These findings indicate that the behaviour of the ants is not merely a matter of maintaining directional consistency or responding directly to sensory input; instead, it should be regarded as the continuation of an internal state ^18^ that drives beacon-aiming.

### Experiment (iii): Decision-making process of beacon-aiming before a land-to-water transition

Finally, to visualise the process of internal state changes, we examined the behaviour of ants before and after transitioning to the adverse condition. In preliminary experiments, we observed that workers of *C. japonicus*, once placed on a small, dry platform surrounded by water (referred to as an ‘island’), spontaneously left the island by swimming after a period of exploration on the platform (which would correspond to reverse aquathlon). Leveraging this behaviour, we investigated whether the ants orient towards a beacon prior to starting to swim. The key point here is whether beacon-aiming occurs before the experience of the adverse substrate condition, i.e., immersion in water, and whether internal state completely switches immediately after a single immersion. Therefore, the behavioural sequence of this experiment was classified into four distinct phases: ‘Initial Phase’: prior to first water contact; ‘Contact Phase’: before first immersion, after first water contact; ‘Immersion Phase’: before initiating swimming, after first immersion; ‘Swimming Phase’: after initiating swimming.

Ants, after released onto an island surrounded by water, showed distinct phases of behaviour. During the Initial Phase, the ants contacted water without a directional bias (Rayleigh test: *R̅* = 0.05, *P* = 0.917, *N* = 33, Figure 4A, B). However, a significant directional bias was evident in the distribution of all water contacts during the Contact Phase (Rayleigh test: *R̅* = 0.27, *P* < 0.001, *N* = 282, mean direction: 23.2°, confidence interval: [6.3°, 39.8°], Figure 4A, C, D, Figure 3A in the supplemental information), which is interpreted as a preference for the right side of the beacon (10°). Similarly, the distribution of the first immersions is significantly biased to the right edge of the beacon (Rayleigh test: *R̅* = 0.34, *P* = 0.033, *N* = 33, mean direction: 6.0°, confidence interval: [−33.5°, 53.0°], Figure 4C), and the distribution of all water immersions is also significantly biased to the right edge of the beacon (Rayleigh test: *R̅* = 0.66, *P* < 0.001, *N* = 33, mean direction: 13.2°, confidence interval: [10.5°, 15.7°], Figure 4A, E,F).

**Figure 4.**
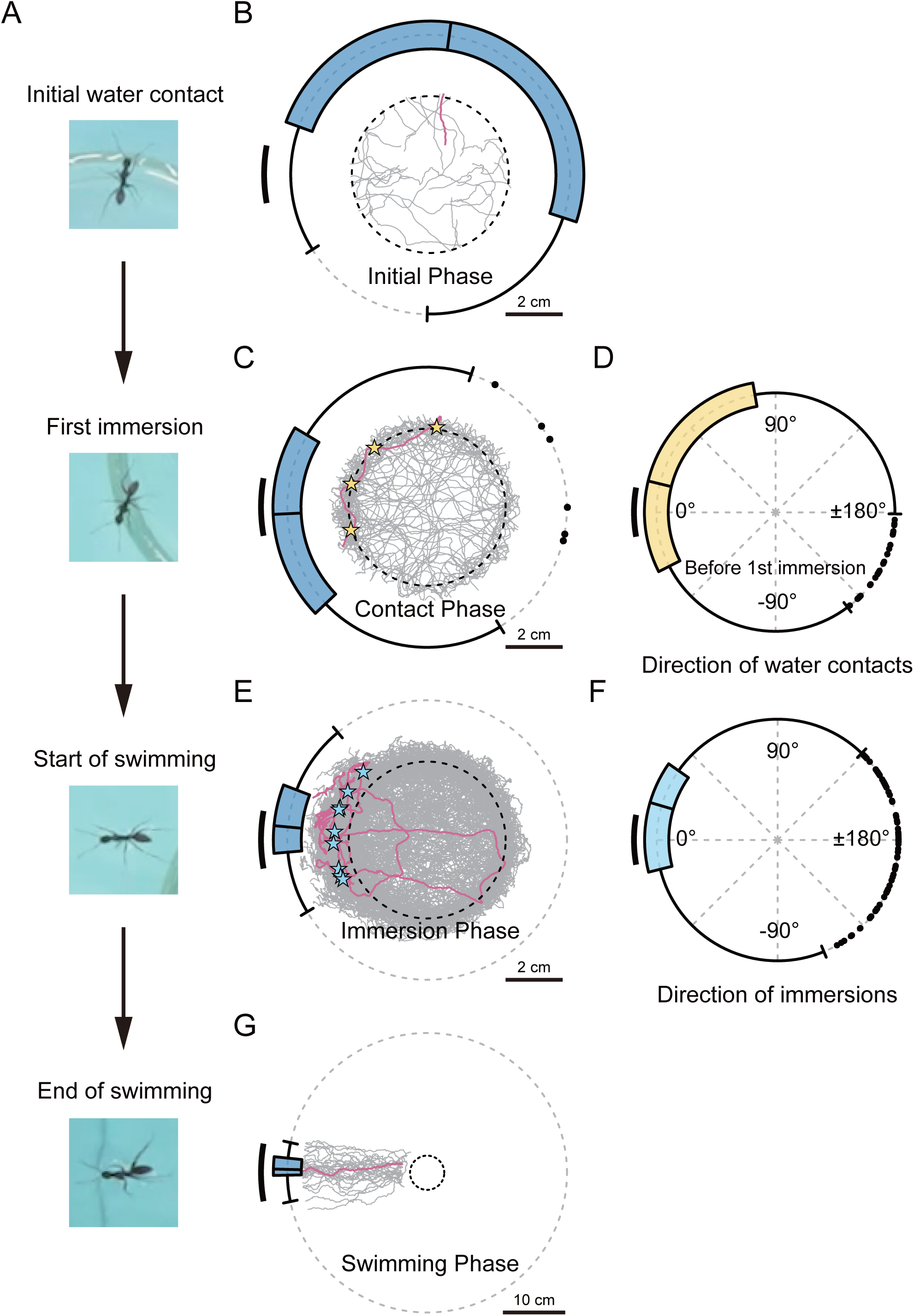
Decision-making process of beacon-aiming. (A) Representative frame images capturing key moments: Initial water contact, First immersion, Start of swimming, and End of swimming. (B-C) Trajectories during the Initial Phase (B) and Contact Phase (C) are displayed along with circular box-and-whisker plots indicating the directions of initial water contacts for all ants (*N* = 33), following the format of Figure 2. A representative trajectory is highlighted in magenta, and the water contact positions during the Contact Phase are marked with light yellow stars. Island edges are shown as black dotted circles, and the trajectory scales are located at the bottom right of each plot. (D) A circular box-and-whisker plot illustrates the directional distribution of all water contacts made by ants before the first immersion, formatted as in Figure 3. (E) Trajectories during the Immersion Phase are presented with circular box-and-whisker plots of initial water contacts, consistent with (B). A representative trajectory (magenta) is included, and positions of all immersions are indicated by light blue stars. (F) A circular box-and-whisker plot shows the distribution of water immersion directions before the ants started swimming, as in (D). (G) Trajectories during the Swimming Phase are depicted with circular box-and-whisker plots showing the final swimming bearings. The format is consistent with (D), and the representative trajectory (magenta) is included.

After several water immersions (Figure 3B in the supplemental information), the ants exhibited ‘pre-swimming behaviour’. Some behaviours classified as immersions were identified as pre-swimming behaviour. This behaviour, characterised by paddling movements while maintaining contact with the shore using at least one hind leg, displayed intermediate characteristics between walking and swimming, complicating quantitative analysis. Most ants began swimming after clear pre-swimming behaviour, showing a directional bias towards the beacon (Rayleigh test: *R̅* = 0.94, *P* < 0.001, *N* = 33, mean direction: 7.0°, confidence interval: [6.1°, 14.8°], Figure 4A, E). During the Swimming Phase, ants finally decided orienting towards the centre of the beacon (Rayleigh test: *R̅* = 0.99, P < 0.001, *N* = 33, mean direction: 1.4°, confidence interval: [−1.0°/3.9°], Figure 4A, G).

These results suggest that, while the ants showed no interest in the beacon during the Initial Phase, they displayed a preference for the sides or edges of the beacon during the Contact Phase, eventually aligning themselves with the centre of the beacon after swimming. Moreover, even after a single immersion, the ants explored various directions (Figure 3B in the supplemental information) while maintaining a bias towards the beacon. These findings indicate that contact with and immersion in water gradually facilitated the formation of an internal state that drives beacon-aiming.

## DISCUSSION

Animals act autonomously in diverse and complex natural environments. The elements defining their behaviour have been classified into 1) reflexes, fixed action patterns, and innate behaviours which can be seen as characteristics of an automatic machine, the extensions of such a machine, or a combination of both, and 2) learning, decision-making, and related processes which are key elements enabling flexible adaptation to diverse natural environments. The primary focus of the research aimed at understanding flexibility and adaptability has been on the latter.

While some ant species are known to orient towards a visual landmark on land ^21,27,29^, recent findings have revealed that *C. japonicus* does not exhibit such orientation under terrestrial conditions ^22^, except in cases where the landmark is associated with appetitive learning ^32^. Our research showed that the attraction to the beacon in this species when crossing water surfaces was triggered by immersion rather than by reception of water itself or the use of swimming-specific locomotor patterns. Changes in orientation decisions depending on substrate conditions are known from a few intertidal species in specific ecological contexts ^33,34^. When forced to walk upside-down under danger of dropping, the ants performed beacon-aiming too. Insects are known to internally monitor their walking movements, detecting reductions in walking speed and/or abnormal leg rotation rates ^35^. Therefore, they are sensitive to the state of contact with the substrate. This is especially true for arboreal arthropods, as dropping from arboreal environments can induce isolation from familiar environments and/or falling onto water surfaces. Unfamiliar environments pose difficulties for navigation, which is a significant problem, particularly for central place foragers ^36,37^. Furthermore, movements across water surfaces pose risks of drowning or predation, especially for small animals ^38–40^. Thus, adverse substrate conditions or even adverse conditions in general are candidates for inducing beacon-aiming. This may apply to other arboreal arthropods or even a multitude of terrestrial animals.

Context-dependent decision-making has been studied in diverse animal species and in various contexts ^41,42^. Although innate behaviours are generally considered to be hard-wired ^10,29^, a certain degree of context-dependence has been reported ^15,43^. However, it remains to be explained whether these context-dependent responses are similar to program behaviours triggered directly by a specific sensory input ^44^ or are instead regulated by transient changes in internal state ^45,46^.

The results of the two paradigms we employed—water-to-land transition (Experiment (ii)) and land-to-water transition (Experiment (iii))—strongly suggest that the innate behaviour of approaching a dark landmark is regulated by internal state in ants (Figure 5A). The former paradigm focuses particularly on persistence of the internal state related to beacon-aiming, while the latter emphasises its emergence. Internal states provide specialized consistency to behavioural tendencies in animals across similar behavioural contexts, such as hunger ^47^, thirst ^48^, social isolation ^49^, and focusing on courtship ^11^. The temporal persistence of internal states has been observed in other arthropods ^50,51^. Meanwhile, behavioural studies that directly observe the real-time dynamics of the gradual changes of internal states remain limited.

**Figure 5.**
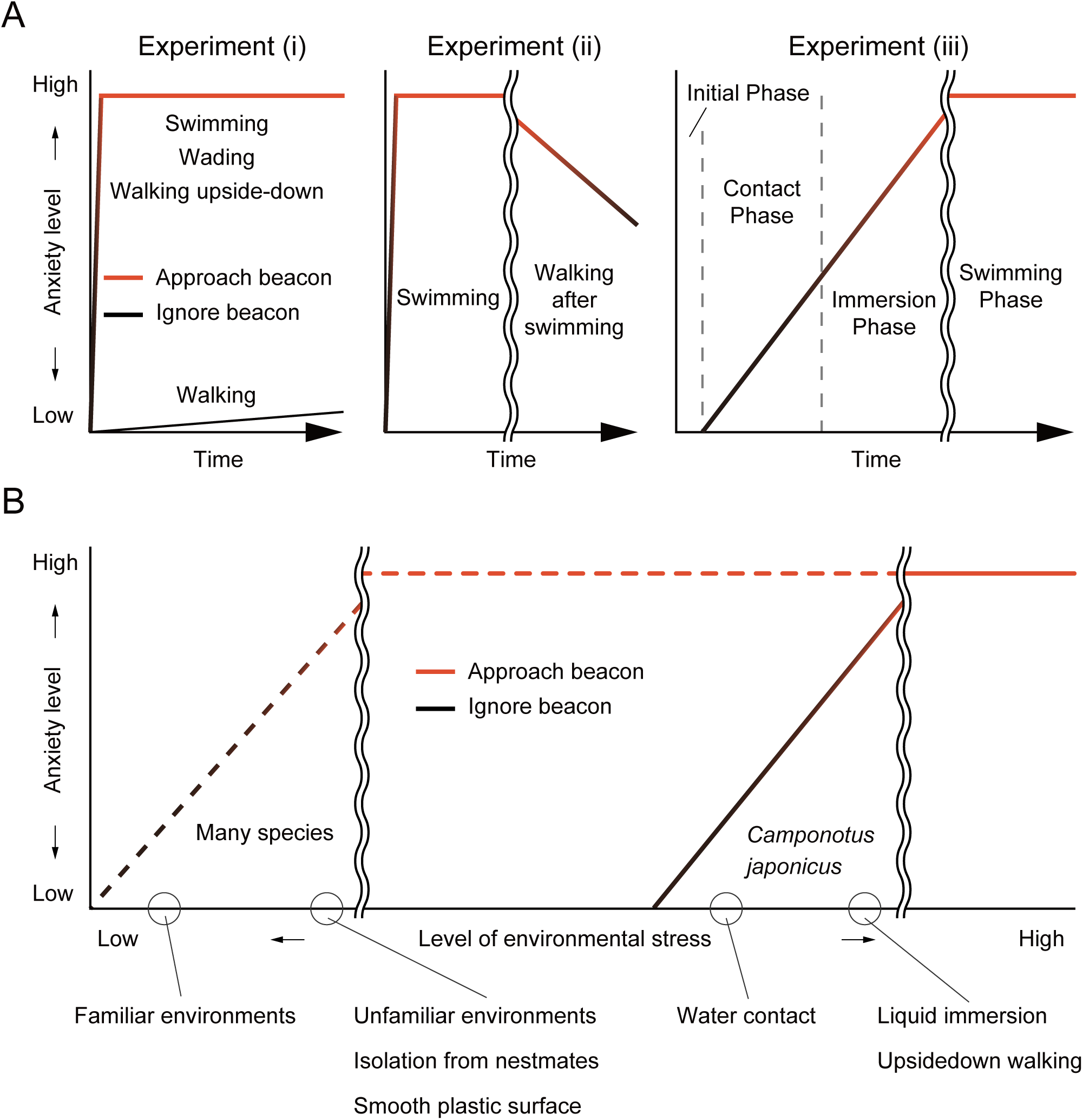
Schematic representations of hypothetical state changes and species-specific responses to environmental stress. (A) Schematic representation of the time series of state changes (anxiety level) in ants under each experimental condition: Single substrate conditions: Walking, Swimming, Wading, and Walking upside-down; Water-to-land transition: Swimming (in the pool) and Walking (after moving onto a dry substrate); Land-to-water transition: Initial Phase, Contact Phase, Immersion Phase, and Swimming Phase. Red line coloration indicates orientation towards the beacon, while black line coloration represents the absence of orientation towards the beacon. (B) A schematic representation based on the hypothesis that interspecific differences in sensitivity to experimental environmental stress, which causes anxiety, determine whether beacon-aiming is present. The dashed line represents many species that exhibit beacon-aiming even under normal substrate conditions, while the solid line represents species that exhibit beacon-aiming only under adverse substrate conditions (*Camponotus japonicus*). Examples of experimental environmental conditions include familiar environments, unfamiliar environments, isolation from nestmates, smooth plastic surfaces, water contact, liquid immersion, and upside-down walking.

As observed in ants introduced onto the island, thorough scanning in unfamiliar environments aligns with previous findings ^52^. The beacon-aiming behaviour of ants on an island was not a binary response but rather a gradually emerging behaviour associated with repeated water contact most likely causing anxiety ^53^. In conclusion, our findings demonstrate that beacon-aiming in the ant *C. japonicus* elicited by adverse substrate conditions can be interpreted as an anxiety-like behaviour.

Anxiety-like behaviour has been studied primarily in rodents ^54^ and fish ^55^, as well as in some arthropods ^56,57^, including insects ^58^, yet detailed research remains scarce for non-model species. The dynamic changes in orientation decisions observed in *C. japonicus* on an island, which might be viewed as a reverse analogue of the Morris water maze, illustrate the process of anxiety-like state formation. To the best of our knowledge, spontaneous escape behaviours from a platform have been studied only in a few invertebrates ^59,60^.

A commonly used experimental paradigm in studies of anxiety is the light-dark test, which induces a preference for darkness (scototaxis) in animals. The simplest explanation for the ecological significance of this preference is that the bright, open areas are innately aversive to many animals, whereas darker spaces are relatively predictable as more secure under predation risks ^61,62^, making them valuable to approach or remain in. Scototaxis has been observed in rodents ^63,64^, fishes ^65^, insects ^66,67^, crustaceans ^68,69^, arachnids ^70^, echinoderms ^71^, and molluscs ^24,72^. Conversely, this innate preference can be altered depending on habitat ^73^, developmental stage ^74^, and time of day ^73^.

The relationship between beacon-aiming and dark preference in general remains to be explained. For instance, tendencies such as preference for an area which includes a number of dark spots over overall darkness ^17^, back-and-forth movement between black stripes in Buridan’s paradigm ^75^, and attraction to a bright beacon against a dark background during swimming ^76^ have been observed. If these behaviours are included in beacon-aiming, the relationship between scototaxis as an anxiety-like behaviour and beacon-aiming would have to be reconsidered.

Is beacon-aiming simply a result of an internal state leading to this behaviour or is there a more sophisticated cognitive process involved to decide to head for the beacon? This is something we cannot decide at this point. Interestingly, beacon-aiming on the island is adapted to handle the future adversity regardless of the actual mechanism that can be the result of a change in internal state ^57^ or cognitive processing ^77^. Examples of responses to future adversity include ants avoiding similar situations by generalizing based on their experiences even after a single emergency encounter ^78^, and grasshoppers that, under chronic predation risk, learn to escape faster and farther ^79^.

Various substances are known to play a major role in changing and maintaining internal states ^50,80^, but knowledge about the mechanisms regulating innate behaviour remains limited ^15,18^. Brain areas known to be involved in beacon-aiming are the central complex, which is necessary for turning during visual following in the context of beacon-aiming ^81^, and the mushroom bodies, responsible for allowing navigation relative to a beacon by learning rather than aiming for it ^12^. The activation of neurons expressing dNPF (*Drosophila* neuropeptide F) ^82^ or octopaminergic neurons ^83^ can induce beacon-aiming towards small objects that *Drosophila* otherwise avoid ^84^. Whether such mechanisms are related to state changes that result in the behaviours we observe around transitions between substrates remains to be seen.

## METHODS

### Animals

Foragers of the Japanese carpenter ant (*Camponotus japonicus* Mayr, 1866) were collected from Komaba I, II campus of the University of Tokyo (35.661°N, 139.684°E; 35.662°N, 139.678°E, respectively) and were individually isolated in 50 ml centrifuge tubes. Prior to the experiments, they were acclimatised under laboratory conditions with light exposure for at least one hour. Only physically intact (no visible damage to legs and antennae) ants were selected for the experiments.

### Experimental procedures

All experiments were conducted inside an opaque cylindrical arena (radius, approximately 30 cm). A vertical black rectangle (beacon) measuring 20° horizontally and 45° vertically from the effective centre of the floor was placed on the inner wall of the arena, based on prior studies ^22,76,84^. To minimise the potential influence of external cues, the beacon was placed in one of three possible positions, each separated by 120°, with trials evenly distributed across these positions. Each trial began with the ant placed at the centre of the floor. The behavioural sequence was recorded using an ILME–FX3 camera (Sony) equipped with a 24–105mm F4 DG OS HSM lens (Sigma) at 59.94 fps, until the ants reached the edge of the circular area used for analysis (arena edge), as defined for each experiment.

In experiment (i), under ‘Land-walking’ conditions, ants were allowed to walk around the arena floor without any obstacles. Meanwhile, ants were forced to swim in the same arena filled with water under ‘Water-swimming’ conditions, and ants were allowed to walk (wade) in shallow water (‘Water-wading’). The behavioural sequences of these conditions were recorded until the ant reached a defined arena edge (radius, 20 cm).

Under ‘Glycerol-swimming’ conditions, ants were forced to swim in a mixture of 80% (v/v) glycerol and 20% (v/v) 2-propanol, which had a dynamic viscosity of 167.1 ± 2.1 cSt (average of three measurements at 24.0°C). Under ‘Tylose-swimming’ conditions, ants were forced to swim across a 1.0% (w/w) tylose (H 10000 P2, SE Tylose, Wiesbaden, Germany) aqueous solution adjusted to closely match the dynamic viscosity of the glycerol-propanol mixture (166.5 ± 1.8 cSt, average of three measurements at 23.7°C). A transparent glass Petri dish (radius, 10 cm), placed at the centre of the arena, was filled with the liquids. The behavioural sequences of these two conditions were recorded until the ant reached a defined arena edge (radius, 9 cm).

Under ‘Upside-down-walking’ conditions, ants were forced to walk upside-down on sandpaper (#2000). The edge of the arena (radius, 9 cm) was defined in alignment with the dimensions of the sandpaper (23 cm × 28 cm).

In experiment (ii), under ‘Beacon Fixed’ conditions, ants were introduced to the pool at the arena centre, and a fixed beacon was presented continuously even after they reached the shore. Meanwhile, under ‘Beacon Displaced’ conditions, the beacon was displaced by 60° immediately upon the ants reaching the shore. Under ‘Walking-without-transition’ conditions, ants were allowed to walk on the dry arena floor without a pool to account for differences in floor height and texture relative to the standard arena used in ‘Land-walking’ conditions. The behavioural sequence was recorded until they reached a defined arena edge (radius, 20 cm), passing through a defined pool edge (radius, 8 cm).

In experiment (iii), a transparent glass cup was inverted to create an island (radius, 2.75 cm) at the centre of the arena, surrounded by water. To prevent ants from inadvertently leaving the island due to their initial momentum, they were first confined within a Fluon-coated cylinder on the island. The cylinder was removed to begin the trial.

The behavioural sequence was classified into four distinct phases:

1. ‘Initial Phase’ (prior to first water contact): This phase lasted until the mandibles crossed the defined edge of the island (radius, 2.75 cm).
2. ‘Contact Phase’ (before first immersion, after first water contact): Starting at the end of the Initial Phase, this phase concluded when the ants’ mandibles crossed the defined immersion edge, represented by a circle with a radius equal to the island’s radius plus two-thirds of the ant’s body length. In this phase, ants contact water mainly with their antennae and forelegs during exploration (defined as water contacts).
3. ‘Immersion Phase’ (before initiating swimming, after first immersion): This phase spanned from the end of the Contact Phase to the moment when the ant lost mechanical contact with the island. In this phase, ants protrude much more over the boundary, apparently exploring beyond for invisible gaps. This is defined as water immersion, but water contacts continue also during this phase.
4. ‘Swimming Phase’: Beginning at the end of the Immersion Phase, this phase ended when the ant reached the defined arena edge (radius, 20 cm).

### Analysis

Trajectories of the ants were obtained using DeepLabCut 2.3.8 ^85^, except for Upside-down-walking, where UMATracker Release-15 ^86^ was instead used. Statistical analyses were conducted as previously described using in Notomi et al (2022 ^22^) Python (https://www.python.org) and the package ‘circular’ (version: 0.4–93) implemented in R ^87^. Circular box-and-whisker plots were generated using a modified version of R code from Buttarazzi et al (2018 ^88^). The Rayleigh test was used to test for uniformity of circular distributions. Differences in circular distributions among experimental groups were evaluated using the Mardia–Watson–Wheeler test. For comparisons made for non-circular data groups, the Mann–Whitney U test was used. All statistical tests were two-tailed and Bonferroni correction applied where necessary.

Further details of Methods are provided in the Document S1 in the supplemental information.

## ACKNOWLEDGEMENTS

We thank all the ants; the members of the Kanzaki and Dobata laboratories for their support, especially Kentaro Matsumura for comments on the manuscript. This research was supported by the Grant-in-Aid for JSPS Fellows (Project Number: 23KJ0665, Y.N.) and Grant-in-Aid for Scientific Research (Project Number: 21H04885, 24H00707, S.D.), funded by the Japan Society for the Promotion of Science (JSPS).

