## supplemental information for "Adaptive decision-making by ants in response to past, imminent, and predicted adversity"

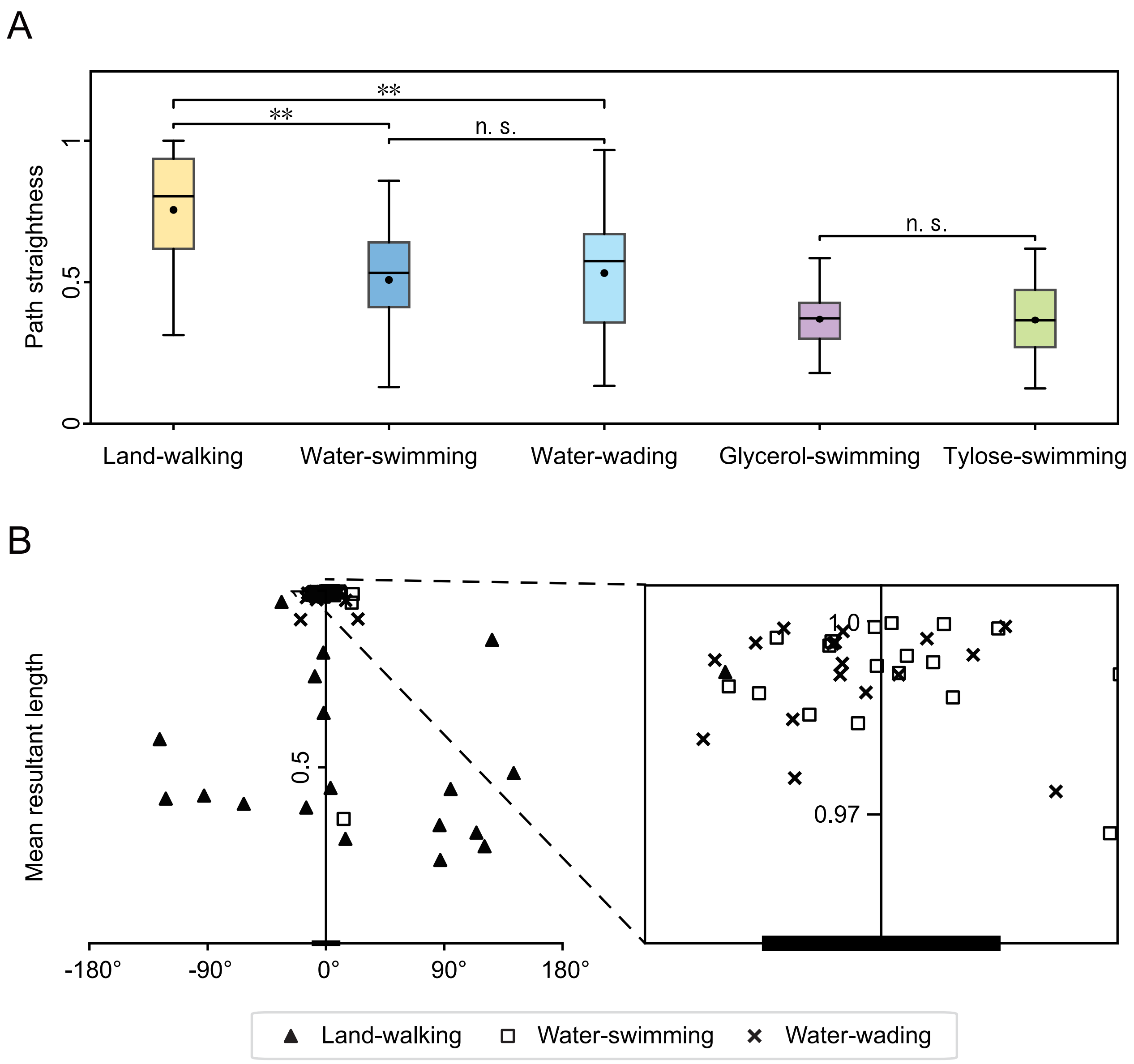


**Figure S1.** (A) Path straightness was analysed across the five conditions: Walking-island; Water-swimming; Water-wading; Glycerol-swimming; Tylose-swimming. ** indicates a significant difference ($P < 0.01$), and n.s. indicates no significant difference ($P > 0.05$), as determined using the Mann–Whitney U test. Box-and-whisker plots illustrate the median, first quartile, third quartile, minimum, maximum, and mean (black-filled circles). Under 'Land-walking' conditions, the path straightness was significantly higher than that under the other two conditions (Mann–Whitney U test: Land-walking and 'Water-swimming': $U = 702$, $P < 0.001$, $N = 57, 57$, respectively; Land-walking and 'Water-wading'; $U = 613$, $P < 0.001$, $N = 57, 57$, respectively), while no significant difference was observed between Water-swimming and Water-wading conditions (Mann–Whitney U test: Water-swimming and Water-wading: $U$ = 1,509, $P$ = 0.516). Additionally, no difference in path straightness was observed between 'Glycerol-swimming' and 'Tylose-swimming' (Mann–Whitney U test: Glycerol-swimming and Tylose-swimming: $U$ = 891, $P$ = 0.897, $N$ = 37, 49, respectively). (B) Scatter plot illustrating the mean resultant length of final bearings for ants across three conditions: Land-walking (filled triangles, $N$ = 19), Water-swimming (open squares, $N$ = 19), and Water-wading (cross marks, $N$ = 19). The MRL quantifies the extent of directional variation across trials for each ant, with values approaching 1 indicating minimal variation and strong directional consistency. When ants consistently aim towards the beacon, the mean direction converges towards 0°. Scatter plots of individual trends reveal that the Water-swimming and Water-wading conditions exhibited minimal variation and a strong concentration around the beacon compared to the Land-walking condition. A black bar indicating the range of the beacon is drawn near the origin in the left figure and under the bottom of the zoomed window in the right figure.

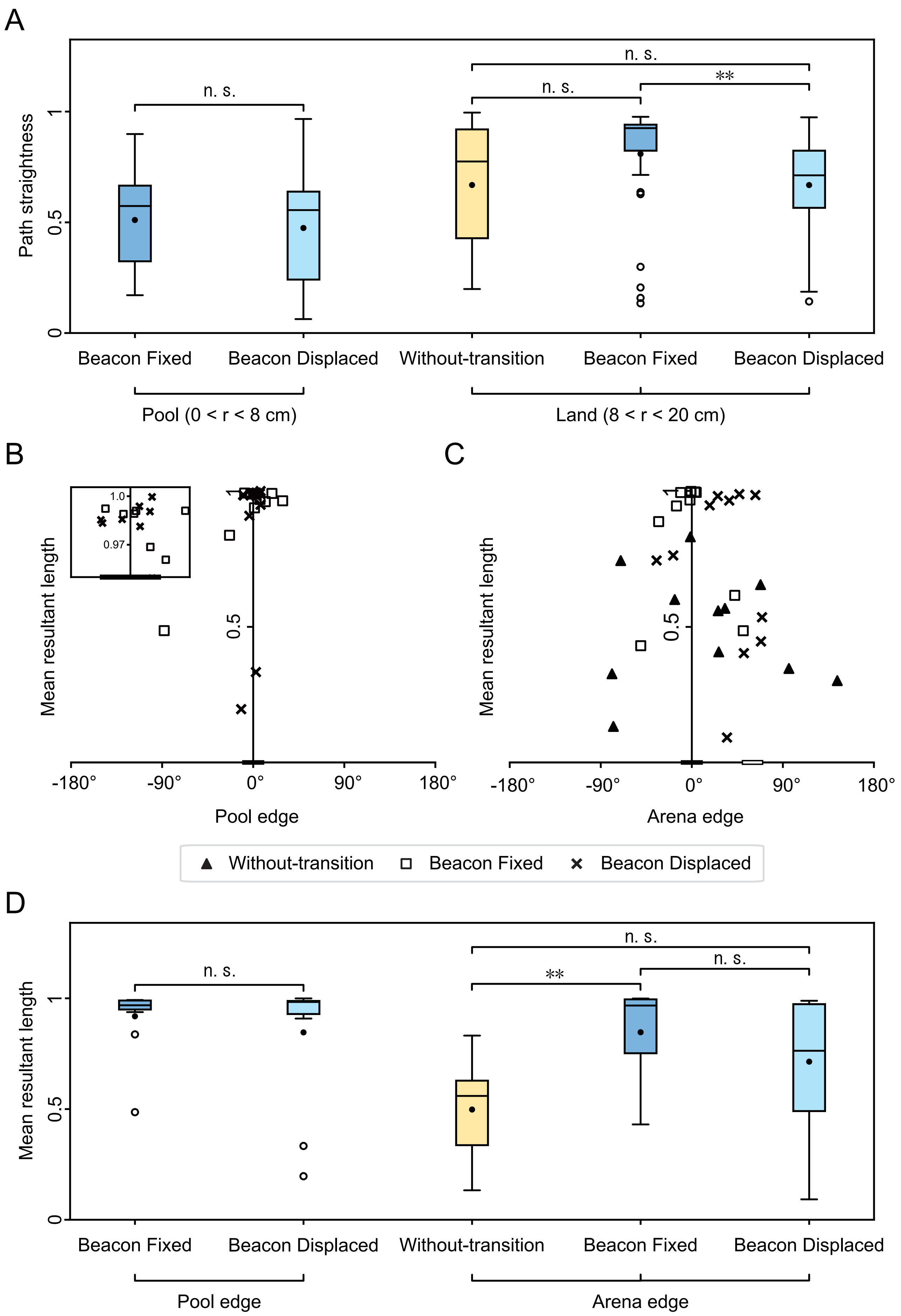
A
Path straightness
n. s.
n. s.
n. s.
**
Beacon Fixed
Beacon Displaced
Without-transition
Beacon Fixed
Beacon Displaced
Pool (0 < r < 8 cm)
Land (8 < r < 20 cm)
B
C
Mean resultant length
1.0
0.97
Pool edge
Arena edge
Without-transition
Beacon Fixed
Beacon Displaced
D
Mean resultant length
n. s.
n. s.
**
n. s.
Pool edge
Arena edge

**Figure S2.** (A) Path straightness during swimming (Pool) and walking (Land) was analysed across three conditions: Walking-without-water ($N = 33$), Beacon Fixed ($N = 33$), and Beacon Displaced ($N = 33$). ** indicates a significant difference ($P < 0.01$), and n.s. indicates no significant difference ($P > 0.05$), as determined using the Mann–Whitney U test. During swimming, there was no significant difference in the path straightness between the Beacon Fixed and Beacon Displaced conditions (Mann–Whitney U test: $U = 490$, $P = 0.491$, electronic supplementary material, figures S2a). During walking, there was no significant difference in the path straightness between Walking-without-transition and Beacon Fixed conditions (Mann–Whitney U test: $U = 438$, $P = 0.175$), and the Beacon Fixed and Beacon Displaced conditions (Mann–Whitney U test: $U = 410$, $P = 0.086$), whereas a significant difference was observed between Beacon Fixed and Beacon Displaced condition (Mann–Whitney U test: $U = 329$, $P = 0.005$). Box-and-whisker plots depict the median, first quartile, third quartile, minimum, maximum, mean (black-filled circles), and outliers, following the format of electronic supplementary material, Figures S1a. (B-C) Scatter plots display the mean resultant length of final bearings at the pool edge (B) and arena edge (C) for ants across the three conditions: Walking-without-water (filled triangles), Beacon Fixed (open squares), and Beacon Displaced (cross marks). The black bar near the origin represents the range of the beacon, and the white bar at 60° represents the range of the Displaced beacon. (D) The mean resultant length at the pool edge and arena edge for all ants across the three conditions (Walking-without-water, Beacon Fixed, and Beacon Displaced) is summarised. Box-and-whisker plots are presented as described in (A). At the pool edge, the swimming final bearings across all ants are not significantly different between Beacon Fixed and Beacon Displaced conditions (Mann–Whitney U test: $U = 59$, $P = 0.332$). At the arena edge, the walking final bearings across all ants are significantly different under Walking-without-transition and Beacon Fixed conditions (Mann–Whitney U test: $U = 25$, $P = 0.002$) but not significantly different between Walking-without-transition and Beacon Displaced conditions (Mann–Whitney U test: $U = 32$, $P = 0.065$) and Beacon Fixed and Beacon Displaced conditions (Mann–Whitney U test: $U = 410$, $P = 0.086$).

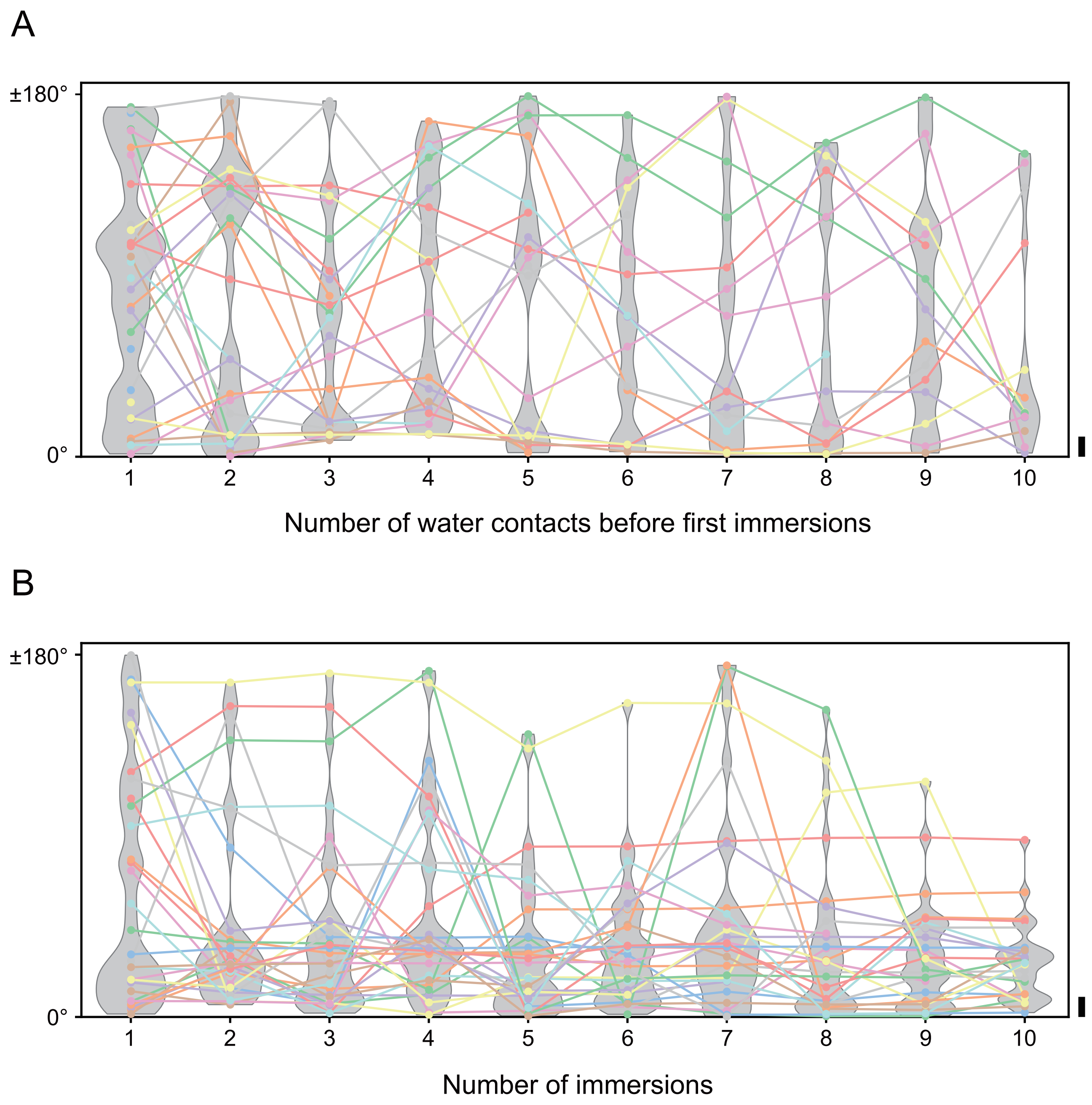


**Figure S3.** (A) Distribution of directions for each water contact before the first immersion (only first 10 contacts shown). The direction of contacts in each trial is represented by filled circles, with consecutive contacts within the same trial connected by lines and shown in the same colour. The violin plots in grey are placed in the background to illustrate the distribution trends. The beacon range is represented as a black vertical on the right side of the plot. (B) Directions at immersion were plotted in the same manner as in (A), only first 10 contacts shown.

**Document S1.**

**a) Experiment (i): influence of substrate conditions and locomotor patterns on beacon-aiming**

The conditions ‘Land-walking’, ‘Water-swimming’, and ‘Water-wading’ were all conducted within the same light blue bucket. The bucket had a base diameter of approximately 58 cm, a top diameter of 70 cm, and a height of 33 cm. The beacon consisted of a vertical black paper strip (width, 10cm; height, 29 cm) attached to the wall of the bucket, positioned such that it subtended 20° horizontally and 45° vertically from the centre of the arena. The position of the beacon was defined in a clockwise direction as viewed from above the arena and was set at 0°, 120°, or 240° for each trial.

The water level in ‘Water-swimming’ was set to a depth of at least 1 cm to allow smooth swimming. The ‘Water-wading’ condition was achieved by adding just enough water to cover the bottom surface, which allowed the ants’ legs to touch the floor.

The arena was illuminated with white LEDs, and the laboratory temperature was maintained at 20–25°C. All experiments were recorded at 59.94 frames per second (fps) using a video camera (ILME–FX3 Cinema Line camera, Sony, Tokyo, Japan with a 24–105mm F4 DG OS HSM lens, Sigma, Kawasaki, Japan) mounted above the arena. Ants were kept in a handheld 50 ml tube and released to the centre of the arena.

The test ants experienced all of the three conditions ‘Land-walking’, ‘Water-swimming', and 'Water-wading'. The ants were divided into two groups, with one group tested in the sequence 'Land-walking' → 'Water-wading' → 'Water-swimming' ($n$ = 9 ants), and the other group tested in the sequence 'Water-wading' → 'Water-swimming' → 'Land-walking' ($n$ = 10 ants). For each condition, the same 19 ants were tested with all three beacon positions, resulting in each ant being tested for 19 × 3 trials. Each ant was given an interval of at least one hour between trials.

**b) Experiment (i): influence of water reception on beacon-aiming**

The experimental arena consisted of an acrylic cylinder (radius, 30 cm; height, 40 cm), covered with white paper on the outside and illuminated with white LEDs. The rectangular beacon (width, 10cm; height, 30 cm) was made of a black paper which was placed on the inner wall in one of three positions as defined in electronic supplementary material, S1, so that it was positioned to subtend 20° horizontally and 45° vertically from the centre of the arena.

A transparent glass Petri dish with a diameter of 20 cm and a height of 2 cm was placed at the centre of the arena and filled to a depth of approximately 1 cm with either an 80% (v/v) glycerol and 20% (v/v) 2-propanol mixture ('Glycerol-swimming') , or a 1.0% (w/w) tylose aqueous solution ('Tylose-swimming') which was adjusted to the same dynamic viscosity as the glycerol mixture.

The chamber used to keep the ants was a lidless 50 ml centrifuge tube, with its inner walls coated with a fluoropolymer (Fluon® PTFE-AGC) by drying an aqueous fluoropolymer dispersion to ensure a smooth descent for the ants. The fluoropolymer coating extended uniformly to the edge of the chamber.

Each ant was tested in a single trial, and the beacon positions were arranged in three directions as described in (a), with approximately equal numbers of trials applied ('Glycerol-swimming': 0°: $n$ = 13 ants, 120°: $n$ = 15 ants, 240°: $n$ = 19 ants; 'Tylose-swimming': 0°: $n$ = 15 ants, 120°: $n$ = 17 ants, 240°: $n$ = 20 ants). Ants that did not reach the arena edge within 5 minutes were excluded.

**c) Experiment (i): upside-down walking**

The experimental arena consisted of an acrylic cylinder (radius, 30 cm; height, 40 cm). A white acrylic board with sandpaper (#2000, DCCS-1P from Fuji Star, Japan) affixed to its bottom side and with a central hole (radius, 0.4 cm) was placed atop the cylinder. The arena was illuminated from below using white LEDs, and the entire setup was positioned on a transparent acrylic board to allow video recording from below. The rectangular beacon (width, 10cm; height, 30 cm) was made of black paper and placed on the inner wall at one of three positions, as described in (a). From the centre of the arena, the beacon subtended 20° horizontally and 45° vertically.

To ensure ants could smoothly initiate upside-down walking, they were introduced via a chamber positioned above the white acrylic sheet. The inner walls of the chamber were fluoropolymer-coated. Ants were introduced through an upper hole in the chamber, and they voluntarily transitioned to the underside of the white acrylic board (i.e. onto the sandpaper surface). Video recording was conducted until the ants reached the arena edge. All ants moved to the sandpaper within 5 minutes from the chamber. Each ant was tested in a single trial, and the beacon positions were arranged in three directions as described in (a).

**d) Experiment (ii): Persistence of beacon-aiming decision made after water-to-land transition**

All conditions were conducted in the same light blue bucket as used in experiment (1). An inset composed of a light blue extruded polystyrene foam disc with a diameter of 42 cm was placed on the floor of the arena (figure 1b). This disc was constructed by bonding two 2 cm thick boards together using waterproof adhesive. The top board featured a circular hole with a diameter of 16 cm, creating a water-filled pool sealed with silicone compound. The pool's water level was filled to the rim (i.e. depth > 2 cm), ensuring that all ants displayed swimming locomotion patterns.

The experiments were conducted in the following sequence: 'Walking-without-transition' → 'Beacon Fixed' → 'Beacon Displaced'. For each condition, the same 11 ants were tested with all three beacon positions, resulting in each ant being tested for 11 × 3 trials. Each condition was separated by an interval of at least one hour. The same ants were used across all conditions and were kept on a dry substrate during the intervals between trials. The beacon positions were arranged in three directions as described in (a).

**e) Experiment (iii): Decision-making process of beacon-aiming before a land-to-water transition**

A transparent cylindrical glass cup (radius, 3.0 cm) was placed upside down at the centre of the arena, and water was added up to the cup's bottom. The bottom of the inverted cup was slightly curved outward (0.25 cm), and aligning the water level with the highest point of the cup created an effective "island" (radius, 2.75 cm) for the ants (figure 1e). To prevent the cup from floating, it was filled with water. The final water depth in the arena, excluding the cup's surface and 0.25 cm of its rim, allowed the ant's locomotion solely by swimming.

Each trial began by introducing an ant into a translucent, Fluon-coated cylinder with a diameter of 3 cm, open at both ends and positioned at the centre of the island, before promptly removing the cylinder. The ant was retrieved before she could contact the arena wall or the beacon during swimming after leaving the island. Three trials were conducted per ant, with different beacon positions for each trial. Following each trial, the ant was allowed to walk on dry tissue paper for at least one minute to remove water droplets from its bodies. The same 10 ants were tested with all three beacon positions, resulting in each ant being tested for 10 × 3 trials. The beacon positions were arranged in three directions as described in (a).

The moment when all body parts of the ants had completely left the island was visually confirmed in videos frame by frame. Additionally, body length was calculated as the average distance between the anterior end of the head and the posterior end of the abdomen across all frames for each ant.

**f) Additional information for analysis**

The video acquired at 59.94 fps was resampled and converted to 60.00 fps before analysis. Trajectories of the ants were obtained by tracking the anterior position of their heads using DeepLabCut, except for Upside-down-walking, where the quality of the obtained videos made it impossible to reliably identify detailed body parts, and centroids were tracked instead of heads using UMATracker. When the likelihood of estimated each body part position calculated in DeepLabCut fell below 0.90 in each frame, the data from DeepLabCut were considered missing values and estimated by linear interpolation from adjacent frames with sufficiently high likelihood. However, the likelihood threshold was set to 0.99 for the experiment (iii), as a large annotation dataset consisting of 7,595 images was prepared. The accuracy of tracking was validated visually by overlaying the computed trajectory data onto the original videos and by creating labelled images for interpolated frames. To improve tracking accuracy, a total of four key points (anterior head, posterior head, anterior abdomen, and posterior abdomen) were annotated for the data from DeepLabCut, which enabled a detailed analysis of the time course of the body axis vector, defined as a vector originating from the posterior abdomen to the anterior head. The behaviour of each ant was analysed until its mandibles (anterior position of their heads) reached the arena edge defined for each experiment. The frames where the ant entered a blind spot of the introduction chamber immediately after release were excluded from the analysis.

The vector connecting the centre of the arena to the position where the ant reached the arena edge was normalised to a unit circle, and its direction was defined as the final bearing of each trial. Similarly, the final bearings at other defined edges were also measured: the final bearing of swimming in experiment (ii) was defined at the first head contact with the pool edge; the final bearings of the Initial Phase, Contact Phase, Immersion Phase, and Swimming Phase in experiment (iii) were defined at the final head position in each respective phase. Additionally, the body axis direction was defined as the bearing of the intersection point between the extension of the body axis vector (from the posterior abdomen to the anterior head) and the wall (circle). The orientation within the arena was defined in a clockwise direction as viewed from above. The orientation of the ant and the body axis direction were calculated as angles relative to the centre of the beacon, set at 0°.

The resultant vector $\vec{R}$ was calculated by summing the unit vectors $\vec{u_i}$ (cos $\theta_i$, sin $\theta_i$) from the three trials for each ant within a condition, where $\theta_i$ represents the final bearing in trial $i$:

$$\vec{R} = \sum_{i=1}^{n} \vec{u_i}$$

The mean resultant length (MRL) was obtained by normalising the magnitude of the resultant vector $\left|\vec{R}\right|$ by the number of trials $n$:

$$\mathrm{MRL} = \frac{\left|\vec{R}\right|}{n}$$

The mean direction $\bar{\theta}$ was calculated as the angle of the resultant vector:

$$\bar{\theta} = \arctan\frac{\sum_{i=1}^{n} sin\theta_i}{\sum_{i=1}^{n} cos\theta_i}$$

The MRL is a quantitative parameter related to the degree of circular consistency in the mean direction and serves as a measure of individual variability in orientation for this study. An MRL value of 1 corresponds to no variability (i.e., a straight trajectory to the beacon), while an MRL value of 0 corresponds to maximal variability (i.e., a uniform distribution). The 95% confidence interval for the mean direction was calculated using the bootstrap method with 1,000 resamples. Path straightness was defined as the ratio of the length of the final vector to the total trajectory distance.

For experiment (iii), the number of water contact events was counted as the number of times the anterior head crossed the island edge. Similarly, the number of immersion events was defined as the number of times the anterior head crossed the immersion edge.
